# Ephaptic coupling can explain variability in neural activity

**DOI:** 10.64898/2025.12.21.695758

**Authors:** Dimitris A Pinotsis, Earl K Miller

## Abstract

The waxing and waning cortical oscillatory power correlates with function and disease. This cross-trial variability has been thought to be due to neuromodulation, uncertainty encoding, and/or changes in cortical excitability. Here, we report evidence that it is also due to fluctuations in ephaptic influences of mesoscale electric fields. We analyzed LFP data from the PFC recorded during a spatial delay saccade task. We constructed a model that describes the electric field close to the cortical patch given the neural activity that generates it. This revealed that field-to-neuron interactions (ephaptic coupling strength) was stronger that neuron-to-field and it correlated trial by trial changes in oscillatory power. This suggests a form of circular causality where neural activity and extracellular electric fields continuously shape each other. These results further suggest that mesoscale ephaptic effects help drive the formation of memory ensembles, a prediction of the cytoelectric coupling hypothesis.

## Introduction

Recent work has suggested that cross-trial variability in cortical oscillatory power reflects a meaningful neural signal rather than noise^1,2^. It has been shown to reflect cognitive state^3^, development^4^ and behavioral performance^5,6^. It has also been proposed as a diagnostic marker^2^. Alpha and beta cross-trial variability has been associated with better recall^7^ and deletion^8^. Reducing gamma burst variability can result in more robust memory representations^9^. Cross-trial variability can also support the uncertainty representations suggested by probabilistic inference theories of brain function^10–12^. It has been linked to power modulations^13^, inhibitory neurotransmission^14^ and E/I balance^15^. Here, we propose and test a different hypothesis: Variability can also be due to ephaptic effects of electric fields.

We suggest that extracellular potential changes modulate the activity of neuronal ensembles within a cortical patch, which in turn further influences the extracellular potential. This creates a circular causality loop, where neural activity and extracellular dynamics shape each other. Our hypothesis was motivated by the cytoelectric coupling hypothesis. It posits that efficient information processing and maintenance in the brain rests upon the integrity of mesoscale electric fields, which, in turn, influence neuronal activity and the cytoskeleton^16^.

To test this hypothesis, we used a combined model that describes both the extracellular electric field close to the cortical patch (near—field) *and* its source, the local field potentials (LFPs) on the patch that generate the field. We found that the strength of field-LFPs interactions, i.e. the strength of ephaptic coupling, correlates cross-trial variability in oscillatory power. This suggests that ephaptic effects help regulate cross-trial variability in oscillatory power and thus play a role in functions associated with that variability.

## Methods

### Experiment and Data

We re-analyzed LFP data originally reported in^17^ also used in our earlier work^18^. Two adult male monkeys (monkey C, *Macaca fascicularis*, 9 kg; and monkey J, *Macaca mulatta*, 11 kg) were trained to perform an oculomotor spatial delayed response task in accordance with NIH guidelines and approved MIT animal care protocols. On each trial, the animals memorized the location of one of six visual targets (0°, 60°, 120°, 180°, 240°, 300°; 12.5° eccentricity) over a 750-ms delay period and then made a saccade to the remembered location. Correct responses were rewarded with water, and incorrect responses were not.

LFPs were recorded using three 32-electrode chronic arrays implanted unilaterally in FEF (Supplementary Figure 1A). Each array consisted of a 2 × 2 mm grid with 400-μm spacing, and implant sites were determined using structural MRI and anatomical atlases. From each electrode, we recorded threshold-crossing spike waveforms and local field potentials (LFPs), the latter extracted using a fourth-order Butterworth low-pass filter (500-Hz cutoff) and sampled at 1 kHz (Cerebus, Blackrock Microsystems). Analyses focused on LFPs during the delay period, when monkeys held the cued location in memory. We assumed each electrode sampled its local neural population and modeled each cortical area as a set of 32 recording sites with monotonically numbered, spatially adjacent electrodes.

### Near field ephaptic effects on cortical ensembles

To describe ephaptic effects on neural activity, we used a combined model of neural activity and the electric field produced by it. This was introduced in ^18^, where we studied how the brain’s extracellular electric potentials *V ^e^* and fields 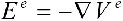 interact with neural activity,*V ^m^*. The membrane potential *V^m^* of a neural ensemble was described using a deep neural field model^19^

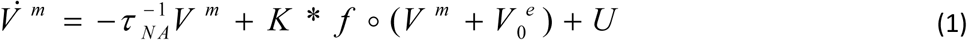

which includes three terms:1) a simple decay, 2) recurrent inputs due to other neurons in the ensemble and diffusive currents resulting from ephaptic coupling and 3) exogenous, stochastic input *U.* In Equation (1), *V ^m^* = *V ^m^* (*z*) depends on the position, *z*, within a cortical patch. *Κ* represents the connectivity or weight matrix that characterizes how the signal undergoes amplification or attenuation as it traverses the neural ensemble. This matrix is shown as colored curves in *Supplementary Figure 1A. V_0_^e^* is the value of the extracellular potential on the surface of the ensemble and *f* refers to the population transfer function ^18^. To sum, Equation (1) describes the temporal evolution of the membrane potential as a result of decay, extrinsic input to the ensemble and diffusion of recurrent input as a result of nearby neuronal activity and ephaptic coupling.

Equation (1) is an example of a neural field model that describes spatially distributed brain responses across extensive brain areas or populations ^20–22^. Neural fields characterise the collective behavior of neural ensembles and the emergence of activity patterns in the brain. They allow us to study activity propagation and interactions across diverse locations. They also incorporate the temporal evolution of these interactions and can be derived from the Wilson Cowan equations ^23^.

The term “deep” reflects the bottleneck architecture of the ReML algorithm used for training^24^. It optimizes a cost function known as the Free Energy, *F*,

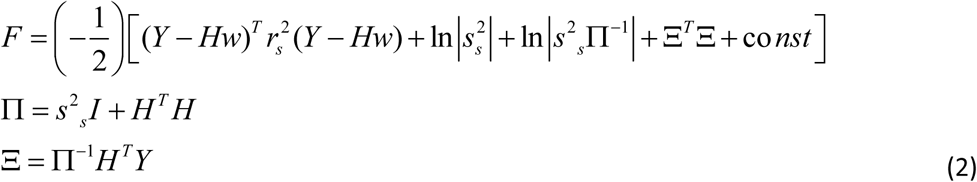

This cost function maximizes the mutual information between the remembered cue and the ensemble activity. After training, one obtains an approximate posterior for the connectivity 𝑟 ∼ 𝑅(𝑤^α^|Ξ, 𝑠_s_^2^𝛱^-1^) with mean 𝛯 and variance 𝑠_s_^2^𝛱^-1^. This is the effective connectivity which in a neural ensemble that has learned to represent a specific stimulus. The principal axes 𝐻^α^, appearing in the 3^rd^ line of Equation (2) are given by 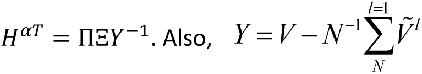 are the mean centered LFP data. They are expressed in terms of the axes and connectivity components via the following Equation, see ^19^ for details.

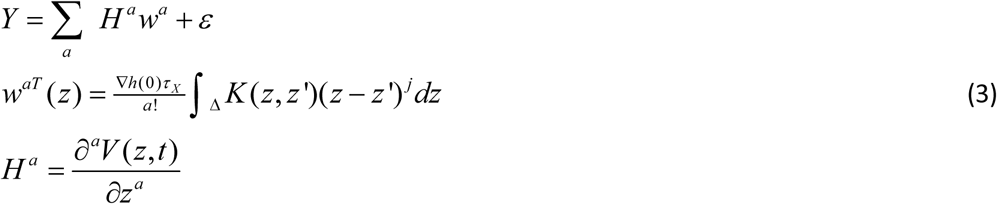

In our previous work, we combined the model given by Equation (2) above, with a model of the electric field (EF; ^18,23^). This model is described below. It allowed us to describe mesoscale fields resulting from neural ensemble activity. This suggested a shift in understanding information maintenance and processing in the brain. Spiking is *not* the end of the story. What needs to be added is a distinct description of the electric field to obtain a combined model of neural activity and electric fields. This combined model then allows one to describe how spiking and electric field levels interact. In other words, spikes are not enough. We need to understand the interplay between neurons and aggregate electric field needs resulting from neurons and the cytoskeleton. Fields might be crucial in organizing neural activity to achieve efficient information processing. We called this hypothesis *cytoelectric coupling* ^16^.

To describe electric fields, we used the *bidomain model* ^23^. This model assumes that the dendrites of cortical pyramidal cells, forming neural ensembles, extend parallelly, allowing for the replacement of their branched dendrites with cylindrical fibers (top panel in Supplementary Figure 1B). This symmetry aligns with the current dipole approximation widely used in human electrophysiology. In this model, pyramidal neurons aligned to produce an EF parallel to apical dendrites, receive synchronous input, generating dipole sources ^25,26^. Electric potential discontinuity, generated by synaptic activity, gives rise to such sources. Extracellular spaces of individual neurons are represented by cylindrical fibers, combined into a unified extracellular space using the principle of superposition (bottom panel in Supplementary Figure 1B). Despite their complex geometry, the parallel alignment of dendrites justifies the cylindrical fiber representation ^27^. This approach aligns with the current dipole approximation commonly used in human electrophysiology ^28^. The potential difference across the membrane boundary creates dipole sources and membrane currents^29^. The model’s symmetry allows the extracellular field and potential to depend on just two spatial variables, *z* and *y,* instead of three. The *z* variable locates positions along the cylinder’s axis, while *y* represents a direction orthogonal to this axis. Below, we derive an expression for the electric field starting from the theory of electromagnetism.

The bidomain model assumes spatial homogeneity and temporal synchrony, like the dipole approximation. While these assumptions may lead to an overestimation of EF, they don’t alter qualitative results, such as ephaptic coupling. The bidomain model furnishes the extracellular potential and field that arises because of the potential discontinuity on either side of the neuron membrane. There, the equation for the temporal evolution of the electric potential to its resting value is given by

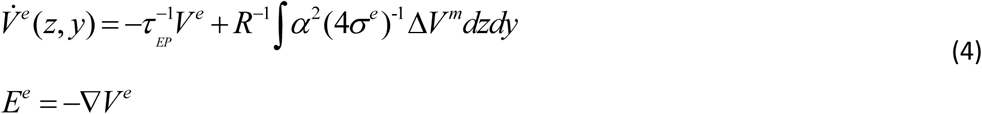

This equation furnishes the relaxation of the electric potential field at a rate 1/*τ_EP_*. The electric potential is generated by charges dispersed along an electrical fiber, parameterized by *z,* arising from the membrane current 1 / *r_i_* Δ *V ^m^*. The elementary source volume element is represented by *dzdy*, with *R* being the distance from the current source to the measurement point of *V ^e^* in the extracellular space. The conductivity *σ ^e^* of the extracellular space and the fiber radius are also included in the extracellular potential. Equation (4) can be written as ^30,31^

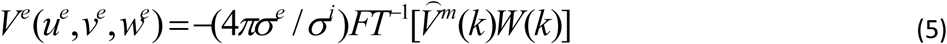

where 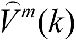 is the Fourier Transform of the transmembrane potential *V^m^* and *FT^-1^* is its inverse Fourier Transform, that is,

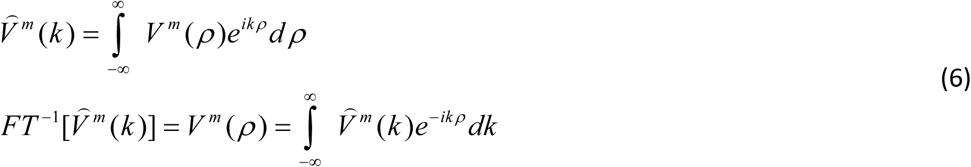

and the function *W*(*k*) is given by

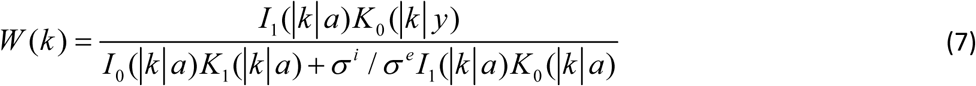

where, *σ^l^*, *l* ={*e*,*i*}, are the extra-and intra-cellular space conductivities and *I*_0_ (*y*),*I*_1_ (*y*), *K*_0_(*y*),*K*_1_(*y*) are modified Bessel functions of the first and second kind ^32^. Equations (5) and (7) are known as the bidomain model of the electric field.

In ^23^, we used the bidomain model above and found that EFs corresponding to neural ensembles exhibited greater stability than neural activity. EFs also had a relatively richer informational content. We posited that this stability allows the brain to compute latent variables underlying the maintenance of the same memory in different trials. In ^18^, we showed that this stability follows from the theory of complex systems and articulated a mathematical framework for ephaptic coupling. We showed that ephaptic coupling is a special case of a general theoretical result known as the *slaving principle* from Complex Systems^18,33^, see *Discussion* for more details.

### Neural activity and electric fields influence each other

Above, we considered ephaptic coupling in relation to the slaving principle. This suggests that electric fields enslave neuronal activity^16,18^. The mathematical expression of this result can be found after expanding the membrane and extracellular potentials *V^m^* and *V ^e^* in terms of Fourier series, that is 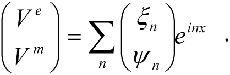 Them, fast relaxing quantities, like the membrane potential modes *ψ _n_*, depend on slowly varying quantities, like the extracellular potential coefficients *ξ _n_*.Substituting the Fourier expansions into Equations (1) and (4), we obtain evolution equations for the Fourier modes (see ^18^ for a proof),

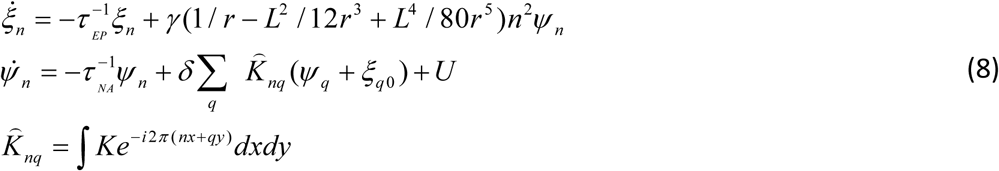

The second line in Equation (8) can be written to describe how the first mode of membrane potential 𝜓_1_ is enslaved by the mode 𝜉_1_ of the extracellular potential --and similarly for higher modes,

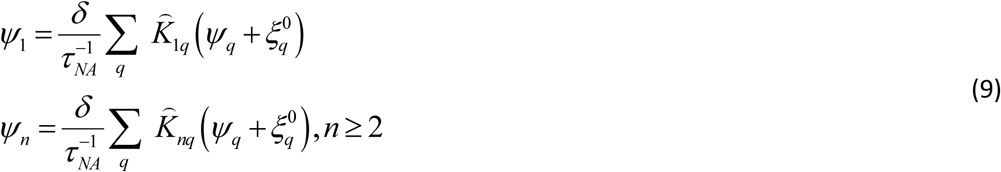

where *q* indexes the modes, 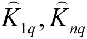 are defined in Equation (8) and *ξ* ^0^ is the boundary value of *ξ* ^0^ on the cylinder of Supplementary Figure 1B. Then, substituting Equation (9) in the first line of Equation (8), we obtain an expression for the first extracellular potential mode 𝜉_1_

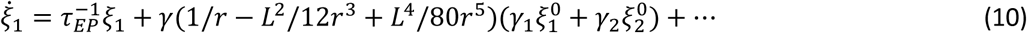

Indeed, from Equation (9) we can express 𝜓_1_ appearing in the first line in Equation (9) by 𝜓_q_. Then *ψ*_1_ = *γ*_1_ *ξ*_1_ ^0^ + *γ*_2_ *ξ*_2_^0^ +… that is, the first mode 𝜓 only depends on near field values, that is, 𝜉_1_^0^, 𝜉_1_^0^,…the electric field very close to the ensemble (the boundary values of the modes 𝜉_1_, 𝜉_2_ on the cylinder of Supplementary Figure 1B). This yields the first membrane potential mode 𝜓_1_ in terms of the terms involving 𝜉1^0^, 𝜉2^0^ etc, the near field values. After substituting this expression into the first line of Equation (8), we obtain Equation (10). This suggests that the evolution (change) of the first extracellular potential mode 𝜉_1_ depends on near field values, that is electric field changes on the boundary of the neural ensemble (𝜉_1_^0^, 𝜉_2_^0^,…).

Assuming some new external input to the neural ensemble when a new memory is represented, the electric field state changes everywhere, including near the membrane. Thus, the extracellular potential mode 𝜉_1_predicted by Equation (10) will change. At the same time, the source, that is the transmembrane potential modes 𝜓_q_, will change simultaneously as predicted by Equation (8). To sum, Equations (8) and (10) predict that perturbing the electric field close to the cortical patch occupied by a neural ensemble changes the extracellular potential *and* its source (the neural activity or LFPs that generate it) – which is circular causality.

### Inter-Trial Variability in Oscillatory Power

We quantified trial-to-trial variability in oscillatory activity by computing both the variance and the coefficient of variation (CV) of spectral LFP power across trials. For each electrode, we first obtained single-trial power using a fast Fourier transform. This yielded a power estimate 𝑃_n_(𝑓) for each trial *n*. Inter-trial power variability was computed as the standard deviation 𝜎_P_(𝑓) = 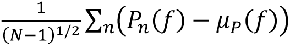 where 𝜇 (𝑓) is the mean power across trials. Because power variance scales with the magnitude of the mean power, we also computed the coefficient of variation 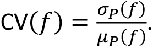

### False Discovery Rate Correction for Correlations

Pairwise Pearson correlations were computed between each single trial LFP power estimate recorded from each electrode (32 locations) and ephaptic coupling strength estimated at each time point (719 samples), yielding 23,008 comparisons per dataset. To control for inflation of Type I error due to multiple comparisons, we applied the Benjamini–Hochberg False Discovery Rate (FDR) procedure^34^ with *q* = 0.05. All *p*-values from the correlation analyses were ranked in ascending order, and the largest *p*-value satisfying the Benjamini–Hochberg criterion was used as the dataset-specific FDR threshold. Correlations with *p*-values below this threshold were considered statistically significant. FDR correction was performed independently for each angle, and only FDR-surviving correlations were carried forward to the group-level summaries and visualizations.

## Results

### Cross-Trial Variability of Oscillatory Power in Data and Model

We first considered cross-trial variability in LFP power recorded during a spatial delay task^17^. In each trial, the angle of a cue presented for 350ms had to be remembered during a fixed delay of 750ms (delayed saccade task; *Supplementary Figure 2*). We analyzed data from the delay period.

Oscillatory LFP power exhibited systematic differences between trials across the delay period and across cued locations. Example responses are shown in Figure 1Ci (single trial LFP spectra averaged across electrodes). We first calculated cross-trial mean and standard deviation. These are shown using blue solid and dashed lines in Figure 1Ai. Standard deviation revealed that oscillatory amplitude fluctuated substantially from trial to trial, indicating that the power of the underlying neural oscillations was not fixed across trials. Mean power was broadly similar across target angles (110.7–118.8), indicating comparable overall oscillatory amplitude during the delay period, see Figure 1Ai). On the other hand, standard deviation showed differences across conditions, with lower variability at 60° (std = 11.46) and 120° (std = 11.02), and higher variability at 180° (std = 13.97), 300° (std = 13.56) and 240° (std = 15.85).

**Figure 1.**
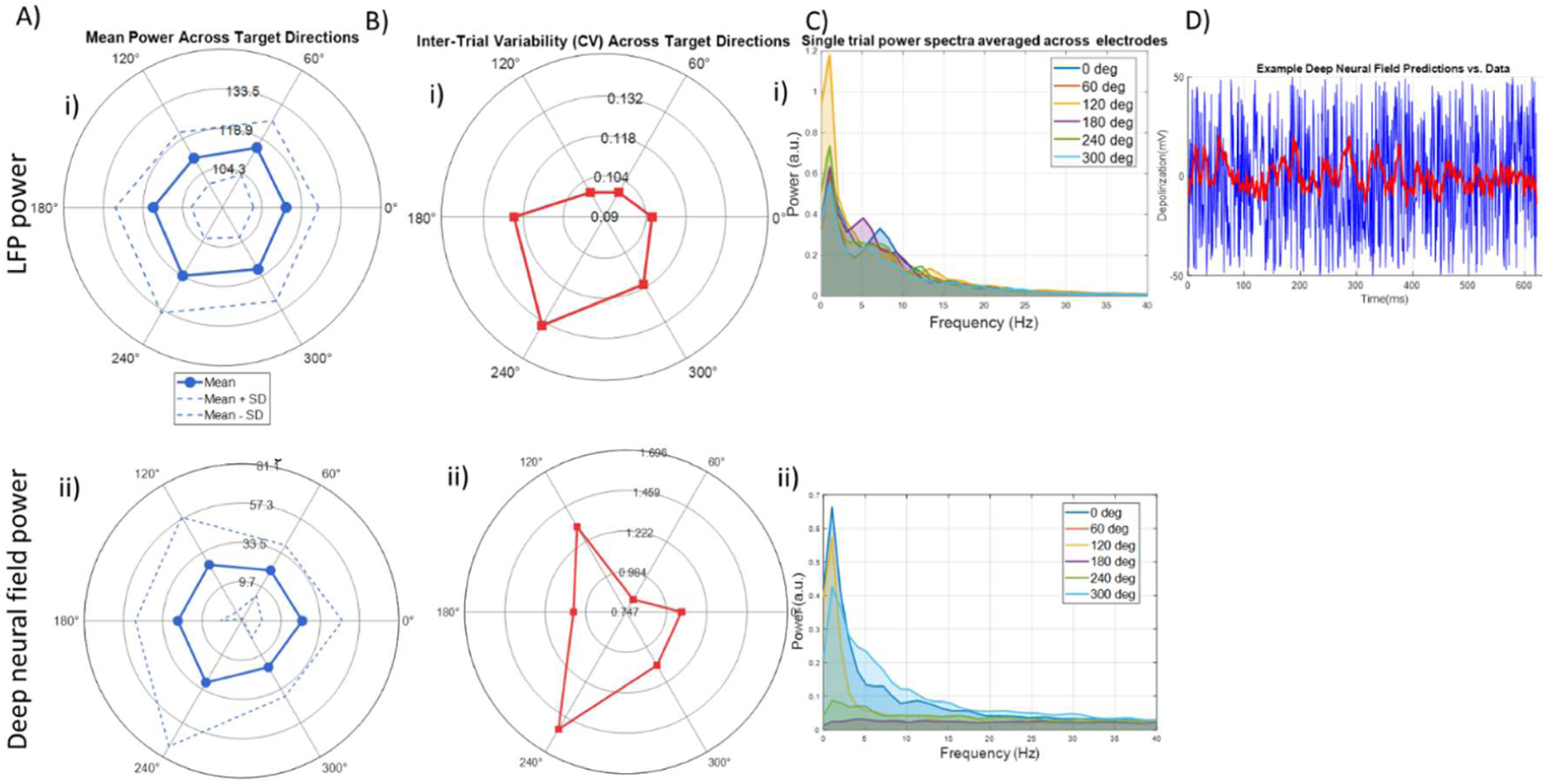
Cross trial variability in LFP responses recorded during a spatial delay task. Top row shows LFP power while bottom row shows the same measures obtained using the predictions of a neural model trained on the same LFP data. A i) Mean LFP power across target directions. ii) Mean deep neural field power. B i) Coefficients of Variation (CV) for LFP responses and ii) deep neural field predictions. C i) Example single trial LFP power spectra averaged across electrodes for different cued angles. ii) Example model predicted power spectra averaged across electrodes. D) Example single trial LFP timeseries (red) and deep neural field model predictions (blue).

The coefficient of variation (CV) confirmed this pattern. CV values were lowest for 60° and 120° (both ≈ 0.10), intermediate for 0° (0.106) and 300° (0.117), and highest for 180° (0.121) and 240° (0.133), see Figure 1Bi). These results indicate that while mean oscillatory power was stable across cued locations, inter-trial variability differed systematically, with 240° and 180° showing the least consistent oscillatory amplitudes across trials.

Below, we study the origin of the cross trial variability in oscillatory power found above. To understand this, we will use a combined neural and electric field model (see *Methods* for details). The neural model predicts neural activity and power spectra for each cued location. Example model predictions are shown in Figure 1D. These are taken from our earlier work on the same dataset, where this model was first used to obtain trial average estimates^18^. In this paper, we focus on single trial estimates.

The model predicts transmembrane depolarization after being trained as an autoencoder using the LFP data considered above. Model predictions and data are shown using blue and red lines in Figure 1D. The model explained about 48-50% of data variance, see also^19^. We called this model a “deep” neural field. The term *deep* reflects the bottleneck architecture of the ReML algorithm used for training, while the term “*neural field*” refers to the general class of models this belongs to, see also *Methods.* The bottom row in Figure 1 includes the same measures discussed earlier based on LFP data. These were not included in^18^. Figure 1Aii) shows the mean power and standard deviations based on predictions of spectral power.

Mean power was broadly similar across target angles (18–29 in a.u.), indicating comparable overall oscillatory amplitude during the delay period. The highest mean power was observed at 240° (29.01), similarly to the LFP data above. Standard deviation was lowest at 60° (std = 17.59) and highest at 240° (44.73), again, like LFP data. CV values were lowest at 60° (0.829) and highest at 240° (1.542), in accord with LFP data too. Intermediate CV values were observed for 0° (1.073), 180° (1.056), and 300° (1.11), and higher for 120° (1.323). Compared with the LFP dataset—which showed relatively low CVs (∼0.10–0.13) and more tightly clustered power values—the model predictions exhibited substantially higher variability overall (CVs ∼0.83–1.54). This can be seen in Figure 1D (blue lines).

Model predictions include much larger trial-to-trial fluctuations and stronger angle-dependent differences in variability. However, the model reproduces the main qualitative findings in the LFP data above: The CV ranking for many of the cued locations is the same. The CV provides a normalized assessment of variability independent of mean power that can be in a different range between the model and data. In other words, as with real LFP data, model predictions of CV values indicate the largest variability at 240° whereas 60° was the most stable. The model also ranked 300° angle, fourth with regard to CV ranking, similarly to LFPs. Results for 0° were also similar, (2^nd^ in the LFP dataset vs 3^rd^ in the model) while results for 120° and 180° were different between the model and the LFP dataset.

Overall, we used CV to describe cross trial variability in oscillatory power and identified cued angles of highest and lowest fluctuations. We also found that a neural model trained with the same LFP data could reproduce this variability for most of the cued angles considered.

### Circular causality between electric fields and LFPs

Above, we found fluctuations in LFP oscillatory power between different remembered angles and characterised them using CV. Next, we asked what drives these fluctuations. Our hypothesis was that they would be driven by the electric field very close to a cortical patch (“near field”) that guides LFPs and also emerges from them, forming a circular causality loop (*Methods*). This also follows from our earlier work^18^, where we had found that trial average causal effects from fields to neurons were much stronger than effects in the reverse direction. In that paper, we had not looked at between trial variability. Here, we focus on single trial estimates of power spectra and ephaptic coupling strength. The precise definition of ephaptic coupling strength is given below after explaining the model used for its derivation.

To compute the ephaptic coupling strength, we considered a model of the electric field, known as the bidomain model^23^ *(Methods)*. This can take LFP data as input and output the corresponding electric fields. These are thought to be generated by the activity of the neural ensemble representing a memory or cued location (near fields recorded with LFP electrodes). When a new trial starts and a new memory (angle) has to be remembered the neural ensemble changes, the effective connectivity on the cortical patch is reconfigured. This changes the near field, which then guides neurons and LFPs into a different state. This, in turn, produces a new field that affects neurons and so on. This is a form of circular causality between neural sources and emerging electric fields (*Methods*). Circular causality suggests that information propagates in both ways between electric fields and neural activity: neural activity generates the electric field and is also controlled by it.

Our hypothesis was that cross trial fluctuations in ephaptic coupling strength reflect oscillatory LFP power variability. To test this, we first computed the single trial estimates of coupling between neural activity and electric fields at the single trial level (ephaptic coupling, EC, strength). We used a variant of Granger Causality (GC), called *spatial GC,* introduced in ^18^. GC is a statistical method traditionally applied to time series data to assess whether knowing the past of one variable (A) improves the prediction of another variable (B) compared to predicting B solely from its own past. If so, it is inferred that information flows from A to B.

To compute GC, a covariance function is first calculated, measuring how strongly a time series is related to itself or to another time series. This requires examining data samples over a certain number of time steps, implying an assumption of finite time, or that information flows at a finite speed from variable A to B. This approach is appropriate when studying processes that occur at measurable speeds over time, such as communication between brain regions, which is where GC has been traditionally applied to ^35–37^. However, because we here consider interactions involving electric fields that occur at very fast speed (about 5000 times faster than synaptic activity^38^), we adapted GC to describe spatial as opposed to temporal correlations and analyzed spatial snapshots of the data at a single time point. We asked if knowing the electric field in one spatial location could improve LFP predictions in a neighbouring, yet unmeasured, location, similar to GC estimates of time-based interactions. The reverse relationship—whether LFPs can predict electric field values—was also tested. The results of the analyses were summarized, and an *F*-test was used to assess the strength of causal interactions between the two variables, following the methodology outlined by ^35^. We called the output of the *F*-test ephaptic coupling (EC) strength. EC strength quantifies the strength of interaction between electric fields and neural activity. We calculated EC strength for all trials and each remembered angle.

These analyses indicated ephaptic effects(Figure 2). Panels 2a)-c) show GC strengths for angles *of 0^0^,60^0^* and *120^0^*. Results for other angles were similar (not shown). Blue violin plots show ephaptic coupling strengths from the field to LFPs (neural activity), while red plots show EC strengths in the reverse direction, from LFPs to the field. These results confirm strong field-to-activity interactions over time, while activity-to-field interactions were less pronounced. The range of blue plots is larger – one order of magnitude larger than the red plots. The spread of the violin plots depicts EC strength fluctuations over time. The variability of top down EC strength (blue) is also comparable across angles, with variances ranging between approximately 5.3 and 6.7.

**Figure 2.**
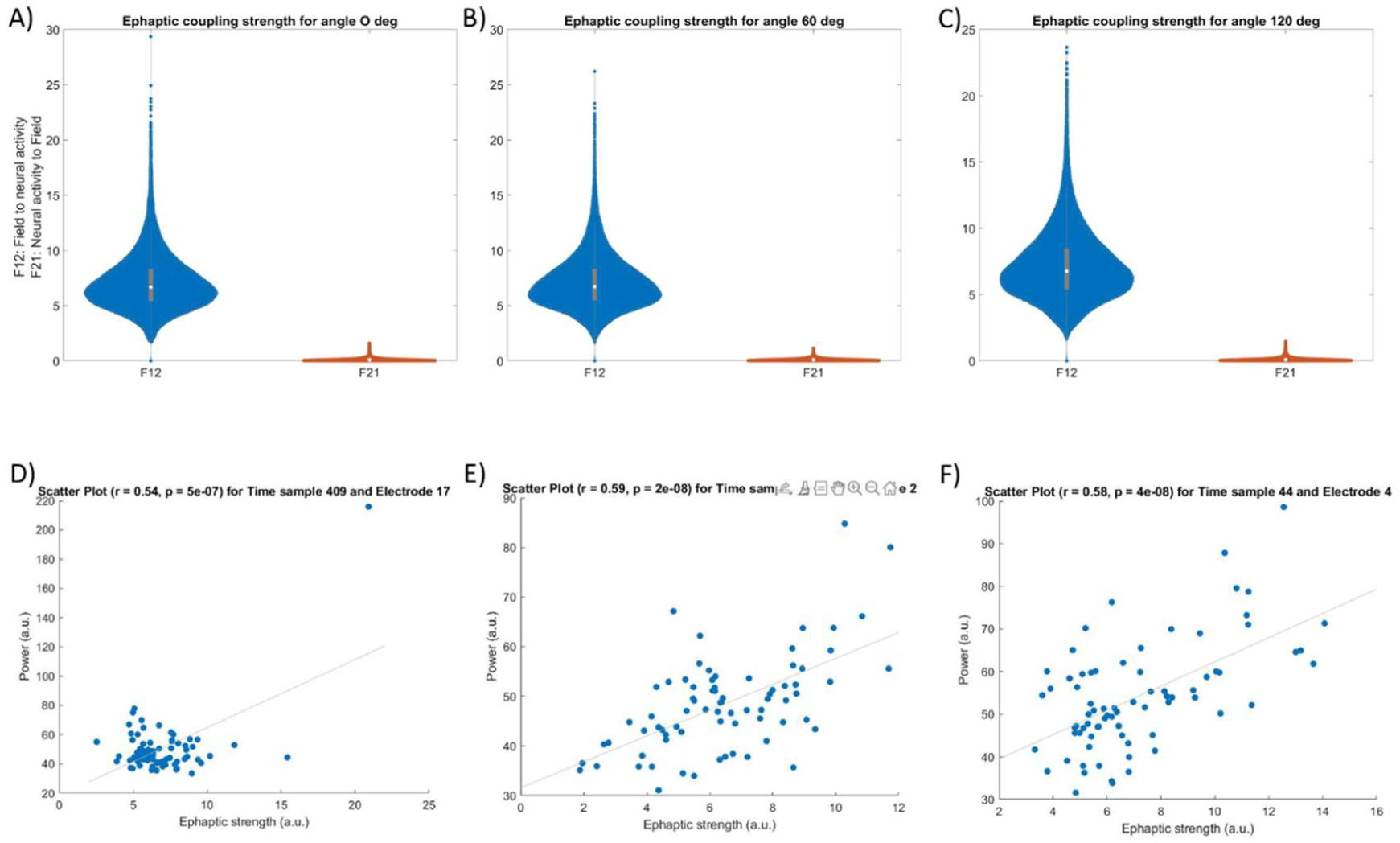
Panels a)-c) show ephaptic coupling strengths for the LFP (neural activity) to electric field direction (red) and the reverse electric field to LFP direction (blue). Strengths were calculated for all angles during each time point during delay Results are shown for angles of 0^0^,60^0^ and 120^0^. Results for other angles are similar (not shown). Panels d)-f) show significant correlations between brain power and ephaptic strength for these three angles at certain time points and locations (electrodes).

To sum, we found that the electric fields that emerge from neural activity, captured with LFPs, turn around and influence this activity in a top down fashion (ephaptic coupling). Information propagated from the mesoscale electric field to LFPs.

### Ephaptic coupling drives oscillatory power fluctuations

Above we found that ephaptic coupling strength varied over time. We also found that neural activity captured with LFPs was guided by electric fields as top down GC strength was larger than in the reverse direction. This suggested that EC strengths and the brain’s oscillatory power might be linked. This is what we studied next. We asked if single trial EC strength samples correlate with LFP oscillatory power recorded from different electrodes (locations) on the multielectrode array. If they do, this suggests that the electric field can drive power fluctuations observed when the brain maintains different memories (angles).

To test this, we computed the correlations between power and EC strength for each remembered angle. Pairwise correlations were computed between the 32 electrode locations and the 719 time samples across 75 trials per remembered angle, yielding 23,008 tests per dataset. Multiple comparisons were controlled using the Benjamini–Hochberg False Discovery Rate (FDR) procedure at *q* = 0.05. The number of FDR-significant correlations varied substantially across the six cued locations.

No correlations survived FDR correction in for angles of 0^0^, 120^0^, or 300^0^, indicating no detectable linear associations that exceeded the FDR threshold in these datasets. In contrast, 60^0^ yielded 1 FDR-significant correlation. The strongest effect occurred between the time point 409 and electrode at location 17 (*r* = 0.544, *p* = 4.54 × 10⁻⁷). More robust FDR-corrected effects were observed for angles of 180^0^ and 240^0^. 180^0^ exhibited 17 significant correlations, with the strongest association between time point 341 and electrode 2 (*r* = 0.594, *p* = 1.97 × 10⁻⁸; FDR threshold *p* < 3.62 × 10⁻⁵). 240^0^ showed 22 significant correlations, with the strongest effect between time point 44 and electrode 4 (*r* = 0.584, *p* = 3.9 × 10⁻⁸; FDR threshold *p* < 4.59 × 10⁻⁵). Overall, FDR-corrected correlations between LFP power and ephaptic coupling strength were present for three angles (60^0^, 180^0^ and 240^0^). The angle of 240^0^ showed the most extensive patterns of association. Examples of significant correlations are shown in Figures 2d)-f). Interestingly, 240^0^ also showed the largest cross trial power variability (see Figure 1). This could relate to homeostasis. We will come back to it in the Discussion.

We then considered predicted spectral power from the deep neural field model considered above (bottom row panels in Figure 1). We wanted to see if model predictions would coincide with our results based on LFPs above. We also wanted to test if the model could provide any further insights in the structure of the above correlations. Similarly to our earlier analyses, pairwise correlations were computed between EC strength and LFP power predicted by the deep neural field model (*Methods*) for each remembered angle. Multiple comparisons were controlled using the Benjamini–Hochberg False Discovery Rate (FDR) procedure at *q* = 0.05.

All six datasets showed substantial numbers of FDR-significant correlations, with the number of surviving associations ranging from 158 to 454 depending on angle. At 0°, 213 correlations survived FDR correction, with the strongest association observed between time point and electrode 27 (*r* = 0.679, *p* = 2.17 × 10⁻¹¹; FDR threshold *p* < 4.59 × 10⁻⁴). At 60°, 188 correlations survived, with the strongest effect between time point 491 and electrode 28 (*r* = 0.673, *p* = 3.69 × 10⁻¹¹; threshold *p* < 3.99 × 10⁻⁴). The 120° angle exhibited 306 significant correlations—the highest among the lower-angle conditions—with a peak correlation between time point 702 and electrode 8 (*r* = 0.656, *p* = 1.73 × 10⁻¹⁰; threshold *p* < 6.58 × 10⁻⁴). At 180°, 158 correlations survived FDR correction, with the strongest association between time point 423 and electrode (*r* = 0.652, *p* = 2.42 × 10⁻¹⁰; threshold *p* < 3.36 × 10⁻⁴). The last two angle conditions showed the most extensive patterns of association: 240° yielded 454 FDR-significant correlations, with a maximal effect between time point 719 and electrode 6 (*r* = 0.642, *p* = 5.37 × 10⁻¹⁰; threshold *p* < 9.84 × 10⁻⁴), and 300° produced 406 significant correlations, with the strongest association between time point 389 and electrode 18 (*r* = 0.677, *p* = 2.53 × 10⁻¹¹; threshold *p* < 8.75 × 10⁻⁴). Overall, all angular conditions exhibited robust FDR-corrected correlations, with the 240° and 300° conditions showing the highest number of significant associations.

To visualize the structure of significant associations in time and space, FDR-surviving correlations for each angular condition were plotted as heatmaps, with time samples along the vertical axis and electrodes along the horizontal axis. These heatmaps reveal clear and systematic patterns across datasets. Fewer correlations survived when using LFP data (Figures 3A and 3B) compared to the deep neural field model to compute the power (Figures 3C and 3D). This can be attributed to low SNR in our recordings and volume conduction. However, results were qualitatively the same.

**Figure 3.**
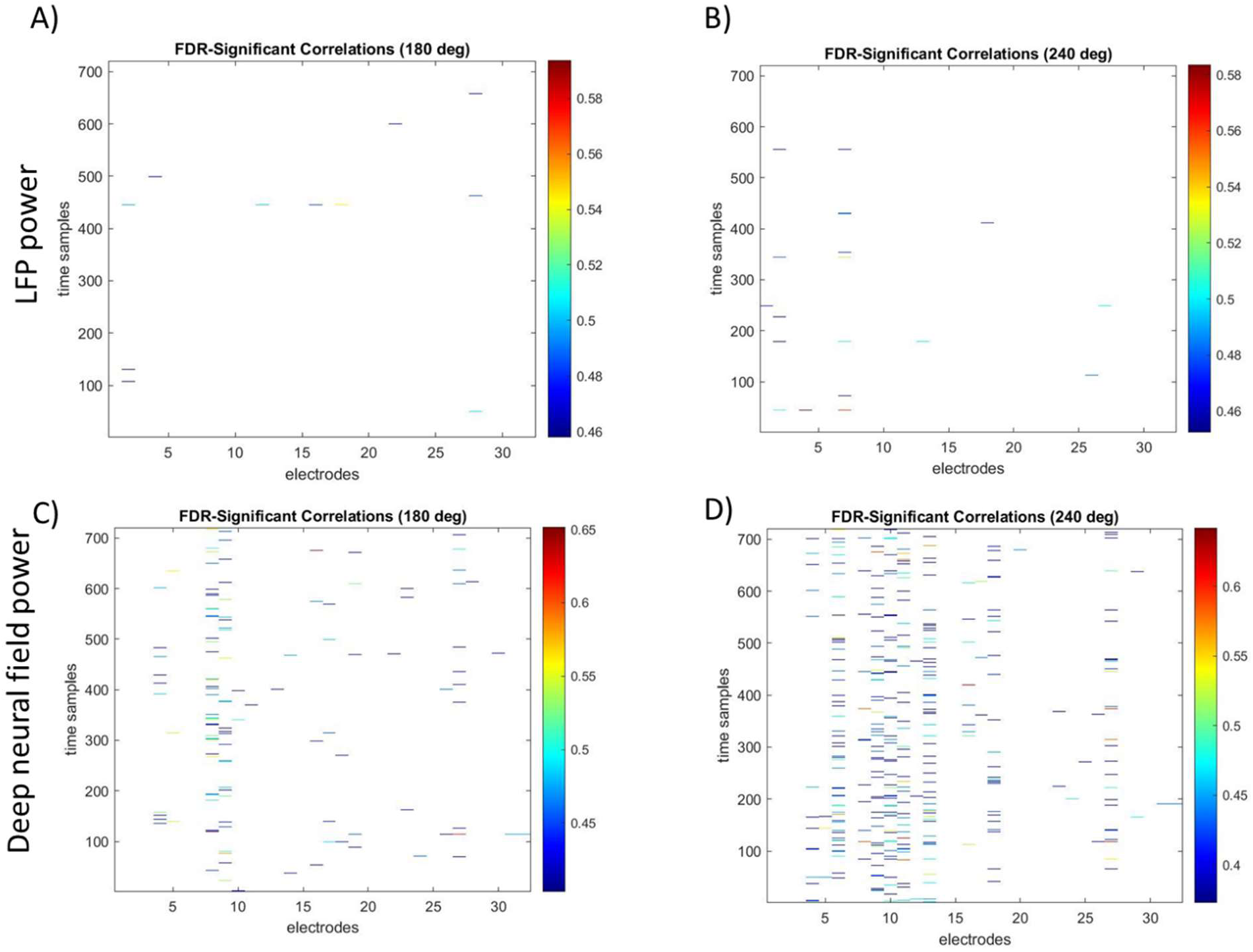
Heat maps of FDR-significant correlations between all time samples and electrodes for cued locations at 180^0^ (panels A,C) and 240^0^ (panels B,D). Panels A) and B) show the results obtained using LFP spectral power, while panels C) and D) show the results obtained using predicted LFP power from the deep neural field model. Significant correlations appeared in spatially localized clusters, indicating that ephaptic effects would affect neurons captured by nearby electrodes at the same time (depicted by small horizontal lines), see text for details.

Significant correlations appeared in spatially localized clusters, indicating that ephaptic effects would affect neurons captured by nearby electrodes at the same time, as expected. These are depicted by small horizontal lines in Figure 3. Also, 240^0^ showed the most extensive patterns of association relative to other angles, in both LFP data and model predictions (panels B,D).

To sum, brain power and ephaptic strength correlated for half of the cued locations when considering LFP power and all locations when considering predicted LFP power from our deep neural field model. We also found that during each trial, information propagated from the mesoscale electric field to LFPs much more than in the reverse direction. During certain epochs of this delay period, the strength of information propagation correlated with oscillatory power at certain locations on the cortical patch. These results support our hypothesis that the electric field sculpts and fine tunes neural activity with high spatial precision and at certain particular moments – giving perhaps rise to oscillatory bursts^39,40^.

## Discussion

We integrated electrophysiological, theoretical, and mesoscale perspectives to reveal how the ephaptic effects from electric fields can contribute to variations in cortical oscillatory power. We found that information propagated from mesoscale electric fields to neural activity, captured with LFPs. The strength of these interactions correlated with variations in oscillatory power over space and time. Furthermore, it seemed that the larger the cross-trial variability in LFP spectral power, the more extensive pattern of such correlations was. In other words, the larger the variability the more evident the top down, organizing effects – a process reminiscent of homeostasis. This follows from the cytoelectric coupling hypothesis^16^. The goal of homeostasis is efficient information processing and memory storage. Cytoelectric coupling suggests how this could be achieved: The electric field oscillations “tunes” the cytoskeleton for efficient information processing, organizing the underlying the spiking activity that help generate the LFP oscillations. The resulting mesoscale organization of neural activity can thus guide the formation of neural ensembles that form memories^18,41,42^.

Ephaptic effects –from electric fields and their transformations to and from potential and metabolic energy through processes reviewed in ^16^—seem to be an application of a general physical principle. This is borrowed from physics^33^ and biology^43^. It suggests that ensemble formation is the result of electric fields that act in consort with inhibitory influences, plasticity and neuromodulation. Electric fields would help form rich-club neural ensembles^44^ and stabilize dynamics by redirecting flow along low-dimensional manifolds embedded in the larger phase space. To put it briefly, ensembles would begin as disordered neuronal clusters that are then brought together by top down ephaptic effects.

The top down causality from electric fields to LFPs we found here is consistent with our earlier work ^18,45^. There, we found evidence in support of this hypothesis after considering cross-trial correlations and trial averages of Granger Causality strengths. We showed that electric fields were more stable and evolved more slowly than LFPs, see also^46^. Decoding accuracy for fields was higher than that for LFPs^45^. In^18^, we also quantified the top down information propagation from the electric field to LFPs using Granger Causality. We found robust top down GC interactions, one order of magnitude larger than in the reverse direction and showed that mathematically top down effects follow from the slaving principle of the theory of Synergetics^33^.

The emerging picture is that electric fields serve as control parameters in the theory of Synergetics. Control parameters evolve more slowly and constrain order parameters and enslaved components. Here, order parameters are the latent states, like effective connectivity components ^19^ and enslaved parts are spiking neurons ^39,47^. This is consistent with our observation that the electric field evolves more slowly than the underlying LFPs. Varying a control parameter, such as the electric field, induces various spatial patterns of neural responses, like those observed in cognitive maps ^48^. This also explains the similarity of dynamic patterns of functional connectivity when the same memory is formed ^49^ and the separation of timescales between fields and neural activity that we and others have found^45,50^. The same prediction follows from experiments testing ephaptic coupling ^51^. Also, it agrees with a common assumption in bio-electromagnetism is that the EF bis quasi-static; the tissue reactance is assumed to be negligible and electromagnetic propagation effects can be ignored ^50^. In the language of complex systems and synergetics, it is a parameter that controls (enslaves) neurons (see also *Methods*). It evolves at a much slower timescale than the enslaved neural activity, similarly to Wiener’s “virtual governors”^52^.

Together, these findings suggest that ephaptic coupling operates as a mesoscale organizing principle linking cellular, electrical, and network dynamics. Electric fields, by acting as slow-evolving control parameters, constrain the high-dimensional trajectories of neural activity and guide the self-organization of ensembles. Through their interaction with inhibition, plasticity, and neuromodulation, these fields stabilize functional connectivity patterns and promote memory formation. In this view, memory engrams emerge not solely from synaptic modification, but from the joint influence of electrical self-organization and dynamic field–neuron reciprocity that governs the brain dynamics.

## Supplementary Figures

**Supplementary Figure 1.**
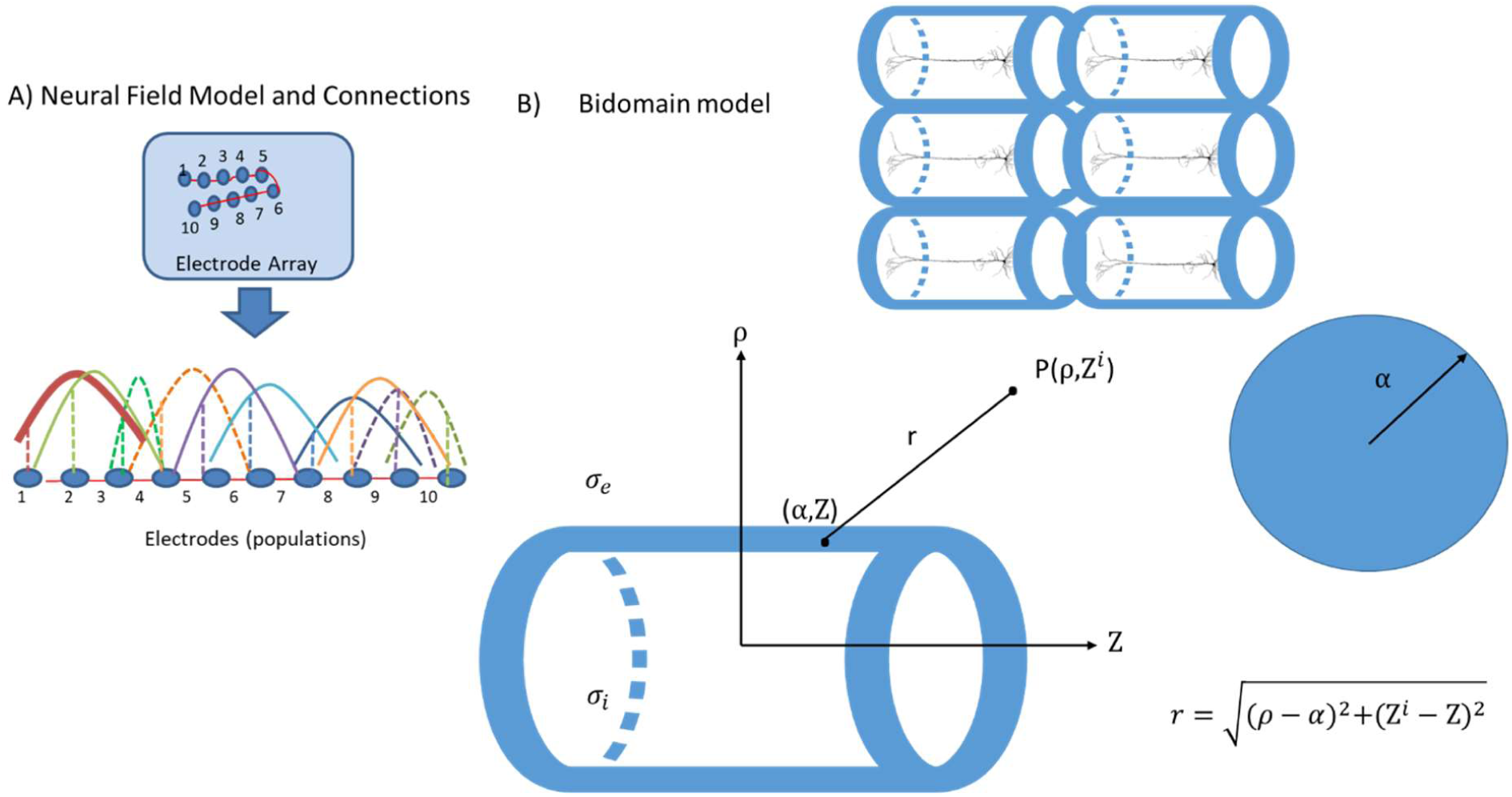

**Supplementary Figure 2.**
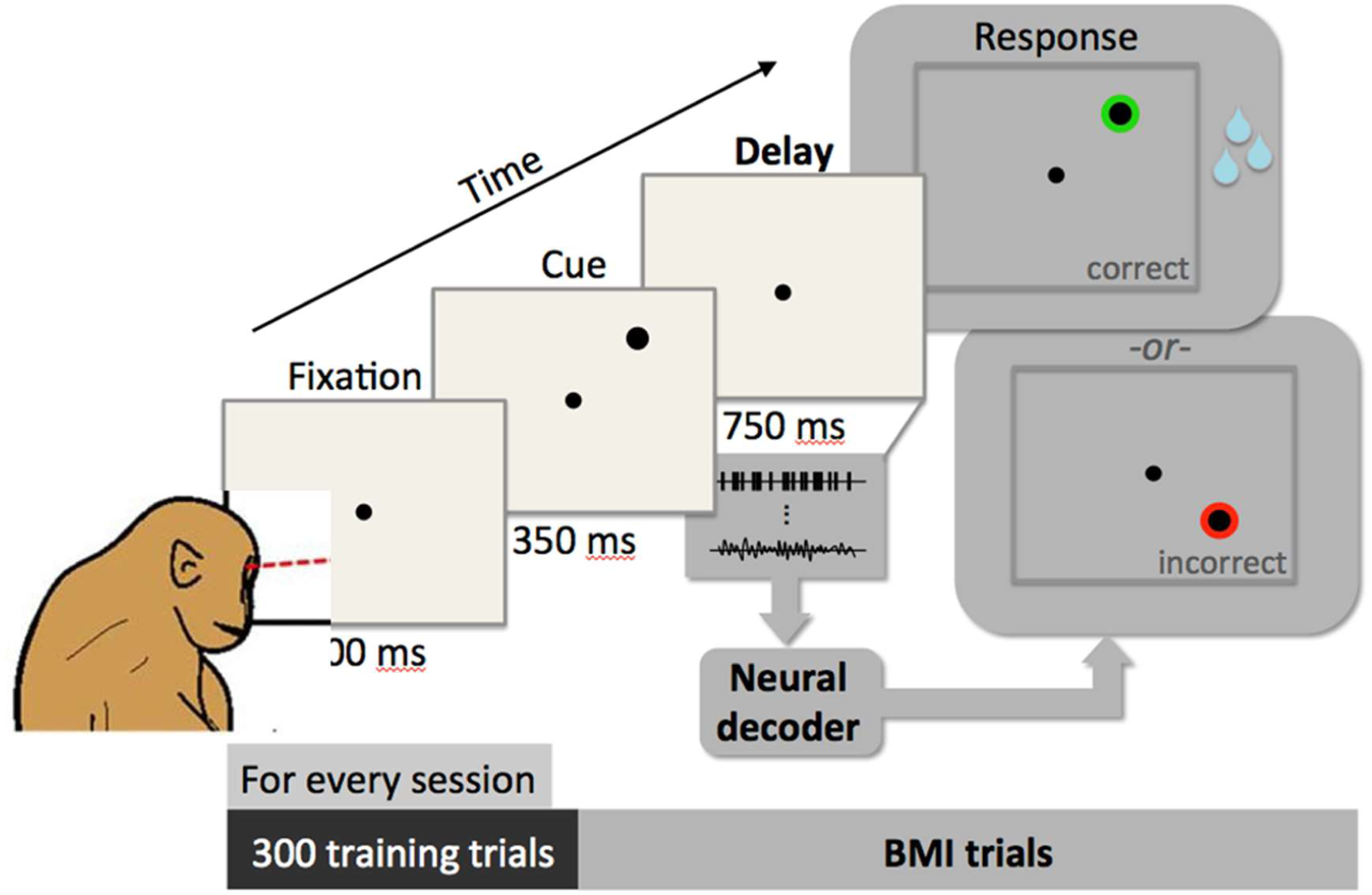

